# Episignatures stratifying ADNP syndrome show modest correlation with phenotype

**DOI:** 10.1101/2020.04.01.014902

**Authors:** Michael S. Breen, Paras Garg, Lara Tang, Danielle Mendonca, Tess Levy, Mafalda Barbosa, Anne B Arnett, Evangeline Kurtz-Nelson, Emanuele Agolini, Agatino Battaglia, Andreas G Chiocchetti, Christine M Freitag, Alicia Garcia-Alcon, Paola Grammatico, Irva Hertz-Picciotto, Yunin Ludena-Rodriguez, Carmen Moreno, Antonio Novelli, Mara Parellada, Giulia Pascolini, Flora Tassone, Dorothy E Grice, Raphael A Bernier, Alexander Kolevzon, Andrew Sharp, Joseph D Buxbaum, Paige M Siper, Silvia De Rubeis

**Affiliations:** Seaver Autism Center for Research and Treatment, Icahn School of Medicine at Mount Sinai, New York, NY 10029, USA; Department of Psychiatry, Icahn School of Medicine at Mount Sinai, New York, NY 10029, USA; Department of Genetics and Genomic Sciences, Icahn School of Medicine at Mount Sinai, New York, NY 10029, USA; The Mindich Child Health and Development Institute, Icahn School of Medicine at Mount Sinai, New York, NY 10029, USA; Graduate School of Biomedical Sciences, Icahn School of Medicine at Mount Sinai, New York, NY 10029, USA; Department of Psychiatry & Behavioral Sciences, University of Washington, Seattle, WA 98195, USA; Laboratory of Medical Genetics Unit, Bambino Gesù Children’s Hospital, Rome, Italy; Department of Developmental Neuroscience, IRCCS “Stella Maris Foundation”, Pisa, Italy; Department of Child and Adolescent Psychiatry, Psychosomatics and Psychotherapy, Autism Therapy and Research Centre of Excellence, University Hospital Frankfurt Goethe University, Deutschordenstr. 50, 60528, Frankfurt am Main, Germany; Child and Adolescent Psychiatry Department, Hospital General Universitario Gregorio Marañón, School of Medicine, Universidad Complutense, IiSGM, CIBERSAM, Madrid, Spain; Medical Genetics, Department of Molecular Medicine, Sapienza University, San Camillo-Forlanini Hospital, Rome, Italy; MIND Institute, School of Medicine, University of California, Davis, USA; Department of Public Health Sciences, School of Medicine, University of California, Davis, USA; Friedman Brain Institute, Icahn School of Medicine at Mount Sinai, New York, NY 10029, USA; Department of Neuroscience, Icahn School of Medicine at Mount Sinai, New York, NY 10029, USA

**Keywords:** Helsmoortel-van der Aa syndrome, ADNP syndrome, DNA methylation, autism spectrum disorder, intellectual disability

## Abstract

ADNP syndrome, also known as Helsmoortel-van Der Aa syndrome, is a neurodevelopmental condition associated with intellectual disability/developmental delay, autism spectrum disorder, and multiple medical comorbidities. ADNP syndrome is caused by mutations in the activity-dependent neuroprotective protein (ADNP). A recent study identified genome-wide DNA methylation changes in 22 individuals with ADNP syndrome, adding to the group of neurodevelopmental disorders with an epigenetic signature. This methylation signature segregated those with ADNP syndrome into two groups, based on the location of the mutations. Here, we conducted an independent study on 24 individuals with ADNP syndrome and replicated the existence of the two, mutation-dependent ADNP episignatures. To probe whether the two distinct episignatures correlate with clinical outcomes, we used deep behavioral and neurobiological data from two prospective cohorts of individuals with a genetic diagnosis of ADNP syndrome. We found limited phenotypic differences between the two ADNP groups, and no evidence that individuals with more widespread methylation changes are more severely affected. Also, in spite of the methylation changes, we observed no profound alterations in the blood transcriptome of individuals with ADNP syndrome. Our data warrant caution in harnessing methylation signatures in ADNP syndrome as a tool for clinical stratification, at least with regards to behavioral phenotypes.

## Main text

ADNP syndrome, also known as Helsmoortel-van Der Aa syndrome, is an autosomal dominant neurodevelopmental disorder (NDD) caused by *de novo* mutations in the *ADNP* gene (MIM #615873) ^1^. The syndrome was first described in 2014 by Helsmoortel and colleagues in ten individuals ^1^. The clinical presentation included intellectual disability/developmental delay (ID/DD), autism spectrum disorder (ASD), facial dysmorphisms, hypotonia, and congenital heart defects (CHD) ^1^. A recent study collating clinical information from medical records for 78 participants across 16 countries has further expanded the clinical spectrum and demonstrated high prevalence of CHD, visual problems, and gastrointestinal problems ^2^.

The *ADNP* gene encodes a multi-functional protein harboring nine zinc fingers ^3^, a homeobox domain for the binding to the DNA ^3^, a binding site for the heterochromatin protein 1 (HP1) ^4^, and a nuclear localization sequence ^3; 5^. In mouse embryonic stem (ES) cells, ADNP interacts with the chromatin remodeler CHD4 and the chromatin architectural protein HP1 to form a complex called ChAHP ^6; 7^. The ChAHP complex binds to specific DNA regions and represses gene expression by locally rendering chromatin inaccessible ^7^. Additionally, the ChAHP modulates the 3D chromatin architecture by antagonizing chromatin looping at CTCF sites ^6^. Within the ChAHP, ADNP mediates the binding to specific DNA sites ^6; 7^. The genes transcriptionally repressed by the ChAHP in ES cells control cell fate decisions ^7^. In fact, *Adnp* null (*Adnp*^−/−^) ES cells undergo spontaneous differentiation and neuronal progenitors derived from them express mesodermal markers instead of neural markers ^7^. In line with the critical role in cell fate specification, *Adnp*^−/−^ embryos have impaired tissue specification, especially of the neural tube, and die *in utero* ^8; 9^. Heterozygous mice, on the other hand, are viable but show developmental delays and behavioral deficits ^10–13^. Of note, ADNP has been described to have extra-nuclear functions critical for synaptic formation and maturation ^12; 14^.

Three recent studies - two on an overlapping sample of 22 individuals with ADNP syndrome ^15; 16^ and a third on an additional 10 ^17^ individuals - reported a genome-wide DNA methylation signature in the syndrome. This observation adds to the body of evidence showing characteristic patterns of methylation changes in the peripheral blood of individuals with NDDs caused by mutations in chromatin factors, including Kabuki ^15; 18; 19^, Claes-Jensen ^15; 20^, Sotos ^15; 21^, Coffin-Siris ^15; 22^, CHARGE ^15; 19^, and Floating-Harbor syndrome ^15; 23^. Importantly, these epi-signatures are syndrome-specific, and have diagnostic value in guiding the interpretation of genetic variants with unclear pathogenicity ^15; 17; 18; 24^. In addition to the signatures found in genetically-defined NDDs, we recently reported an enrichment of epigenetic changes (epimutations) in individuals diagnosed with NDDs and congenital anomalies ^25^, ASD ^26^, and schizophrenia ^26^, suggesting that sporadic epimutations might contribute to the etiology of these disorders. Unlike other syndromes with a methylation signature ^15; 18–23^, the methylation changes observed in ADNP syndrome clearly stratify mutation carriers into two classes: class I, for individuals with mutations located outside a region between nucleotides 2000 and 2340 of the *ADNP* coding sequence (NM_015339.4), and class II, for individuals with mutations between c.2000 and c.2340, including the recurrent mutation p.Tyr719* ^16^. The two episignatures can be harnessed to predict the syndrome and the specific mutational class ^15–17^ but whether they correlate with clinical outcomes or gene expression changes in blood has not been studied.

To address these questions, we studied 43 individuals with a genetic diagnosis of ADNP syndrome from 4 cohorts: Cohort A, collected as part of the Autism Sequencing Consortium ^27; 28^; Cohort R, collected at the Medical Genetics Laboratory of Ospedale San Camillo-Forlanini ^29^ or Ospedale Pediatrico Bambino Gesù, Rome, Italy; Cohort S, prospectively assessed at the Seaver Autism Center for Research and Treatment, Icahn School of Medicine; and, Cohort W, prospectively assessed at the University of Washington. Mutations in Cohort A were identified through research-based whole-exome sequencing ^27; 28^ and validated by Sanger sequencing. Mutations in Cohort R, S, and W were identified and validated in Clinical Laboratory Improvement Amendments (CLIA)-certified laboratories. Participation was approved by the Institutional Review Boards of participating sites. All caregivers provided informed written consent and assent was obtained when appropriate. We analyzed 24 samples (cohorts A, R, and part of S) for methylation analyses; prospective phenotype data from 30 individuals (S and W) for genotype-phenotype analyses; and, 17 samples (part of cohort S) for RNAseq analyses.

We first performed a genome-wide methylation analysis using Ilumina EPIC 850K methylation arrays on peripheral blood DNA isolated from 24 ADNP participants, 19 unaffected age-matched controls, and 14 unaffected siblings from cohorts S, A and R (**Figure 1A**, **Table S1**, **Supplemental Note**). The 24 ADNP participants split evenly into class I and class II mutations (**Figure 1A**). We first verified correspondence between nominal and inferred sex and age ^30^, and predicted the fraction of CD4^+^ T cells, CD8^+^ T cells, natural killer cells, B lymphocytes, monocytes and granulocytes in the samples ^31^ (**Supplemental Note**).

**Figure 1.**
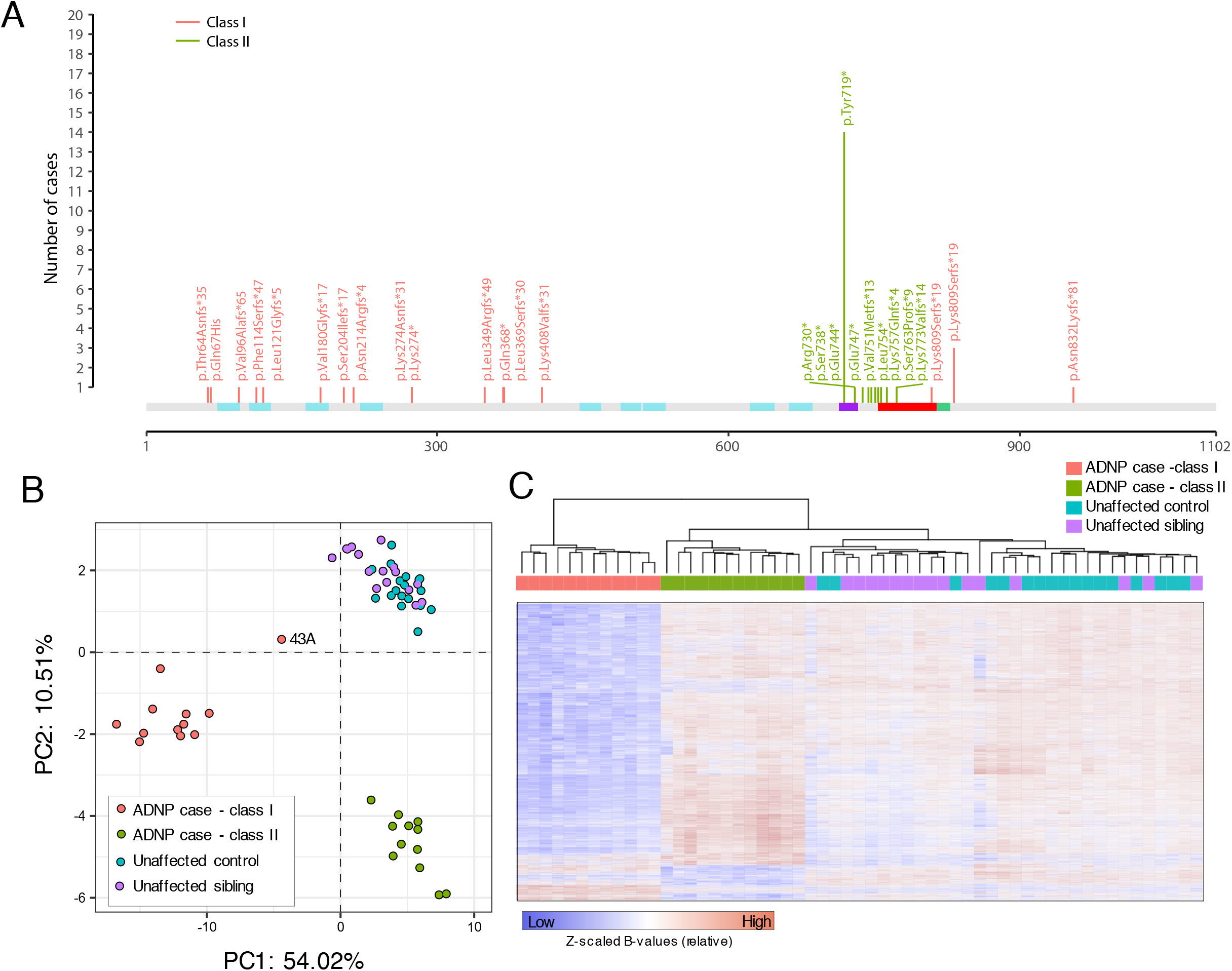
**(A)** Lollipop plot of the *ADNP* mutations in the 42 individuals with point mutations included in the study, including the 24 with methylation data (Individual 1S carrying a deletion is not shown, Table S1). Mutations are described according to the Human Genome Variation Society (HGVS) guidelines for mutation nomenclature. The cDNA and aminoacid positions are annotated according to *ADNP* RefSeq mRNA and protein sequence NM_015339.4 and NP_056154.1. Nucleotide numbering referring to cDNA uses +1 as the A of the ATG translation initiation codon in the reference sequence, with the initiation codon as codon 1. The seven zinc fingers domains (cyan), the nuclear localization sequence ^3; 5^ (purple), the homeobox domain for the binding to the DNA ^3^, and the binding site for the heterochromatin protein 1a (HP1a) ^4^ are shown. Mutations in the c.2157-2317 region are classified as class II and are shown in green; mutations outside of this interval are classified as class I and are depicted in pink. **(B)** Representation of the two principal dimensions in a principal component analysis of the methylation data from 24 ADNP cases (red) 19 unaffected age-matched or 14 unaffected siblings (blue), showing a clear separation between class I cases, class II cases, and controls. **(C)** Hierarchical clustering of 12 class I cases, 12 class II cases, 19 unaffected age-matched or 14 unaffected siblings for the 6,448 autosomal CpGs found differentially methylated in individuals with ADNP mutations belonging to class I.

Subsequently, principal component analysis (PCA) confirmed the clustering of *ADNP* epi-signatures into two groups, according mutation class (**Figure 1B**). These two groups were well separated from unaffected age-matched or sibling controls. Specifically, class II mutations were located between c.2157 and c.2317 of NM_ 015339.4, within the interval defined by Bend and colleagues ^16^. Eight class II participants carry the most recurrent mutation found in *ADNP*, p.Tyr719* (**Figure 1A**, **Table S1**, note that another six p.Tyr719* participants were included in the genotype-phenotype studies). We then performed linear regression using batch, age, sex, predicted blood cell composition and *ADNP* mutation status as independent variables, separately for class I and class II mutations. We selected probes associated with disease status at 1% FDR and with ≥10% mean methylation difference between ADNP mutation carriers and age-matched and unaffected sibling controls (**Supplemental Note**). We identified 6,448 autosomal CpGs that were differentially methylated in class I individuals compared to controls (**Table S2**). Of these differentially methylated sites, 4,143 overlap with the 5,987 previously identified ^16^ (~69%). These sites map to the gene promoter (defined here as ± 2 kb from the transcriptional start sites) and/or gene body (transcription start to transcription end) of 2,802 autosomal Refseq genes, and most hypomethylated in ADNP cases (**Figure 1C, Table S2**). Differentially methylated genes in class I are enriched in neuronal and synaptic genes (“neuron cell-cell adhesion” and “receptor localization to synapse” as top Gene Ontology processes) and in risk genes for ASD ^28^ (26 genes, *P*<0.0001), DD/ID ^32^ (119 genes, *P*=4×10^−4^), and congenital heart defects (CHD) (24 genes, *P*=0.005) (**Supplemental Note**). In line with earlier findings ^16^, class II individuals show a much more modest episignature, with 2,582 differentially methylated autosomal CpGs (**Figure 1C, Table S2**), 1,007 of which overlap with the 1,374 previously described ^16^ (~73%). The 1,442 unique RefSeq genes with altered methylation in the promoter and/or gene body are enriched in DD/ID genes ^32^ (69 genes, *P*=5×10^−4^), ASD ^28^ (11 genes, *P*=0.013) but not CHD (10 genes, *P*=0.13). Interestingly, class I and class II episignatures shared 888 differentially methylated probes (1% FDR and with ≥10% mean methylation difference from controls for both classes). However, the direction of change for these CpGs was inversely correlated, with hypomethylation in class I and hypermethylation in class II (Pearson, *R*=-0.31, *P*=2.2×10^−16^) (**Figure S1**).

Given the striking divergence between the episignatures in class I and II (**Figure 1B**, **1C**, **Figure S1**) and the clustering of class II mutations (**Figure 1A**), we hypothesized that the two types of mutations might differ in their functional impact. Most pathogenic or likely pathogenic mutations in *ADNP* are nonsense or frameshift mutations. Most mutations, including 40 out of the 43 in this study, fall in exon 3, after the last exon-exon junction and are predicted to avoid nonsense-mediated decay (NMD). A previous study has in fact shown that mutant RNAs can be detected in the blood of individuals with ADNP syndrome ^1^, and that mutant ADNP proteins exogenously expressed in HEK293 cells are translated into truncated proteins that can undergo proteasome-mediated degradation or mislocalize in the cytoplasm ^5^. We therefore asked whether class I and class II mutations located in exon 3 differ in NMD escape. To test this hypothesis, we performed Sanger sequencing on the cDNA amplified from RNA isolated from peripheral blood of individuals carrying three class I (p.Phe114Serfs*47, p.Leu349Argfs*49, and pLeu369Serfs*30) and two class II mutations (p.Tyr719* and p.Glu747*) (**Table S1**, **Figure S2A-B**). In all samples examined, we observed biallelic expression of *ADNP*, with both reference and mutant alleles being expressed, confirming that mutant *ADNP* alleles escape NMD (**Figure S2A-B**). We observed no differences between class I and class II mutations, indicating that a differential NMD escape is likely not the mechanism underlying the divergent episignature. We replicated this observation by examining mutation-mapping reads in genome-wide RNA-sequencing (RNA-seq) from 17 ADNP participants (**Table S1**, **Figure S2C-E**). Notably, all mutations had non-zero RNA-seq read coverage and the *ADNP* locus was expressed at low levels (FPKM, 3.03±0.44). When we compared the overall expression of mutant alleles to the expression of reference alleles, we observed lower expression among the mutant reads on average across all samples (*P*-value for difference in log median expression =0.012; **Figure S2E**). Our data, alongside previous evidence ^1^, indicate that the ADNP pathogenic mutations are not leading to classical haploinsufficiency, but rather to the expression of mutant protein which might or might not gain additional functions. Interestingly, one of our ADNP participants (1S) carried a deletion extending from the 5’UTR to the second exon, suggesting that class I mutations are more likely resulting from a loss of function. Also, one class I ADNP participant showed attenuated genome-wide methylation changes (43A, **Figure 1B**). Notably, this individual carries the most distal frameshift mutation (**Table S1**, **Figure 1A**). Similarly, Bend and colleagues found a carrier of a p.Tyr780* mutation having more modest changes compared to other class II mutations ^16^. These independent observations suggest that more terminally truncated ADNP mutant proteins may retain partial function associated with an attenuated epigenetic signature.

Genotype-phenotype correlations have begun to emerge in ADNP syndrome. Previous retrospective review of the medical records of 78 *ADNP* participants reported that carriers of p.Tyr719* have a higher rates of high pain thresholds and walked significantly later than individuals with other types of mutations ^2^. Individuals with p.Tyr719* or adjacent mutations might also have higher risk of blepharophimosis ^29; 33; 34^. An initial analysis of Bend and colleagues on three class I and six class II individuals did not identify differences between the two groups ^16^, although with minimal power. To further probe potential phenotypic differences and relate them to epigenetic changes, we used extensive, prospectively collected clinical data for two cohorts, Cohorts S (class I n=10, class II n=12) and W (n=5/class). Given the profound methylation changes in the blood of individuals with class I mutations, we initially conjectured that individuals with class I mutations would have more severe clinical manifestations. To test this hypothesis, we contrasted data for 70 variables in the cohorts S and W separately and combined (**Table 1, Table S3**). Remarkably, both groups displayed similar levels of intellectual disability, language impairment, ADHD diagnoses, and medical problems. However, we replicated earlier findings ^2^ of a larger delay in first walking independently in class II individuals (class I=22±2.67 months, class II=36.33±6.45 months, *P* <0.0001) (**Table 1**). We also observed some evidence for a higher prevalence and severity of ASD in class II (class I=4/15, class II=12/17, *P* <0.02) (**Table 1**). There was also a trend towards higher rate of self-injurious behavior in class II, but this failed to reach statistical significance (**Table S3**). Importantly, these differences hold up also when we considered class II without the p.Tyr719* mutation (*n*=9), indicating that p.Tyr719* is not acting as an outlier that skews the results. Overall, the striking divergent methylation patterns in class I and class II individuals did not translate into robust phenotypic differences between the two groups, but there is evidence for subtler associations.

**Table 1.** Comparison between clinical measures collected prospectively from 15 class I and 17 class II *ADNP* cases in cohorts S and W. Table S3 presents the analysis of additional measures, broken down by cohort. Shapiro-Wilk test was used to assess normality of all continuous variables and either a one-way analysis of variance (ANOVA) or Wilcoxon rank sum test were implemented accordingly. Chi-squared tests with Yates correction were used to test discrete variables. Abbreviations: NA, data not available.

| **Table 1** | **Class I (n=15)** | **Class II (n=17)** | **P-value** |
| --- | --- | --- | --- |
| Sex | 10 F, 5 M | 5 F, 12 M | 0.080 |
| Age (years) | 7.33 ± 3.33 | 7.18 ± 4.56 | 0.910 |
| Gastrointestinal Problems | 11 | 14 | 0.851 |
| Hypothyroidism | 4 | 2 | 0.587 |
| Structural Brain Changes | 3 | 7 | 0.369 |
| Hypotonia | 11 | 15 | 1.000 |
| Sleep Disturbance | 11 | 12 | 1.000 |
| Obstructive Sleep Apnea | 5 | 1 | 0.185 |
| Congenital Heart Defect | 7 | 5 | 0.522 |
| Early Tooth Eruption | 8 | 7 | 0.388 |
| Seizures | 3 | 7 | 0.364 |
| Vision Problems | 10 | 16 | 0.126 |
| Hearing Problems | 6 | 3 | 0.313 |
| Recurrent Infections | 6 | 7 | 1.000 |
| Feeding Issues | 3 | 9 | 0.120 |
| Gait Abnormalities | 13 | 15 | 0.464 |
| Autism spectrum disorder | 4 | 12 | **0.020** |
| Intellectual disability | 15 | 16 | 1.000 |
| Attention Deficit/Hyperactivity Disorder | 6 | 6 | 1.000 |
| Nonverbal/Minimally Verbal | 9 | 13 | 0.365 |
| Motor Delays | 14 | 17 | 0.949 |
| Sensory Symptoms | 15 | 17 | 1.000 |
| High Pain Threshold | 10 | 15 | 0.252 |
| First Walked Independently | 22 ± 2.67 | 36.33 ± 6.45 | **2.0e-7** |
| First Single Word | 38.71 ± 20.53 | 33.6 ± 12.47 | 0.457 |
| First Phrase | 80 ± 23.51 | 60 ± 15.74 | 0.094 |
| Full Scale IQ/Developmental Quotient (DQ) | 36.23 ± 13.82 | 35.72 ± 17.55 | 0.930 |
| Nonverbal IQ/DQ | 37.68 ± 14.34 | 37.97 ± 19.04 | 0.960 |
| Verbal IQ/DQ | 35.92 ± 14.53 | 34.33 ± 17.97 | 0.800 |
| ADOS-2 Comparison Score | 4.27 ± 1.83 | 6.4 ± 2.32 | **0.010** |
| PPVT-4 (Standard Score) | 60.5 ± 14.7 | 45.75 ± 20.41 | 0.060 |
| EVT-2 (Standard Score) | 53.25 ± 14.9 | 45.14 ± 18.33 | 0.370 |
| Beery VMI-6 (Standard Score) | 58.17 ± 15.99 | 48.7 ± 11.24 | 0.240 |
| Vineland Adaptive Behavior Composite | 53 ± 13.11 | 48.59 ± 12.86 | 0.350 |

We next asked whether changes at the gene expression level could be predicted based on differentially methylated CpGs and also cluster *ADNP* mutation carriers from controls. Principal component analysis was performed on RNA-seq data from 17 ADNP participants and 19 unaffected siblings using genes harboring differentially methylated CpGs within their respective gene body; no distinct stratification was observed between ADNP samples and unaffected siblings (**Figure S3A**). Subsequently, differential gene expression analysis tested for genes that were over- or under-expressed in *ADNP* samples relative to unaffected siblings and identified two genes associated with class I mutations and nine genes associated with class II mutations under an FDR < 0.05, each with small individual effect sizes (**Table S2**). A competitive gene-set ranking approach was used to functionally annotate genome-wide trends in gene expression. First, we note a poor consistency between significantly hypermethylated and hypomethylated CpGs relative to genes that are under-expressed and over-expressed in *ADNP* samples, respectively (**Figure S3B-C**). Second, we performed GO enrichment analysis using our previously identified biological processes found to be significantly enriched for differentially methylated CpGs and confirmed a significant over-expression of genes mapping to gated and ion channel activity as well as passive transmembrane transporter activity (**Figure S3C**). Notably, there was no significant enrichment for ASD, DD/ID or CHD genes, despite a trend of these genes to be over-expressed in ADNP samples. Finally, an exploratory GO enrichment analysis was performed and identified significant under-expression of toll-like receptor signaling in class I ADNP samples. Class II ADNP samples displayed under-expression of neuroactive ligand receptors and extracellular matrix organization and over-expression of protein translation, initiation and termination. (**Table S3D-E**). Overall, despite the critical role of ADNP on chromatin accessibility and architecture ^6; 7^ and the methylation changes found in individuals with pathogenic ADNP mutations, we did not observe profound alterations in corresponding gene expression profiles.

Here we replicated and expanded earlier findings ^15–17^ of DNA methylation changes in ADNP syndrome. The mechanisms leading to these methylation changes are still unknown. While initial observations suggested an association of ADNP with subunits of the chromatin remodeling complex called SWI/SNF ^9^, a recent study failed to replicate this finding ^7^ and found instead that ADNP interacts with CHD4 and HP1 to form a complex (ChAHP) that restricts local chromatin accessibility ^7^ and controls chromatin looping ^6^. Defective ChAHP complexes formed by mutant ADNP might potentially change accessibility for DNA methyltransferases and/or demethylases and explain the methylation changes found here. It is still puzzling, however, why mutations in residues 719-773 (or 719-780 ^15; 16^) have a different effect than all other mutations. In fact, even within the groups of 888 differentially methylated CpGs shared between the two groups, the direction of change is opposite, with more hypomethylation in class I and more hypermethylation in class II. This indicates that the two groups of mutant ADNP proteins affect DNA methylation via different mechanisms. The first important observation is that even frameshift and nonsense mutations do not lead to NMD (**Figure S2** and ^1^), suggesting that these mutations lead to the expression of truncated proteins. These truncated proteins might act as hypomorphic or gain-of-function alleles. The fact that an individual carrying a deletion extending from the 5’UTR to exon 2 clusters with class I mutations (1S) and that the most terminal class I mutation has an attenuated signature (p.Ser955Argfs*36 in 43A) would suggest that this group of mutations might act as hypomorphic alleles. Experiments introducing GFP-tagged ADNP mutants in HEK293T cells have suggested a distinct pattern of expression and localization based on the location of the mutation, dividing the mutations in N-terminal (in the first 412 residues), central (in the 473-719 region) and C-terminal (from residue 719) ^5^. N-terminal mutants were targeted to proteosomal degradation, expression of central mutations was restricted to the cytoplasm, and expression of C-terminal mutants were expressed in the nucleus but showed reduced co-localization with pericentromeric heterochromatin ^5^. These data would suggest that N-terminal and C-terminal mutations reduce protein activity, while mutations in the central domain are more detrimental. However, the methylation data in this and previous studies ^15; 16^ show that mutations in residues 719-780 (and not in residues 473-719) behave differently from all others. Also, independent evidence in ES cells carrying an homozygous p.Tyr718* mutation (corresponding to human p.Tyr719*) show that the mutated ADNP reaches the nucleus ^7^. Interestingly, this mutated ADNP binds to CHD4 and to DNA, despite the loss of the homeobox, but cannot interact with HP1 ^7^. Since HP1 mediates the repression activity of the ChAHP complexes, ADNP target genes are overexpressed in the p.Tyr718* mutant ES cells ^7^. While the analyses from Cappuyins et al. ^5^ and Ostapcuk et al. ^7^ are beginning to explore the functional impact of different ADNP mutations, relevance to the clinical syndrome is complicated by the use of overexpression systems ^5^ or the use of homozygous targeting ^7^. Future studies on patient-derived cells would be important to better dissect the differential impact of the clinical mutations.

Prior analyses on three class I and six class II individuals failed to detect phenotypic differences between the two groups ^16^. Our prospectively collected clinical data on 15 class I and 17 class II individuals show that the two mutational groups differ in the mean age of first walking and rates of ASD (**Table 1**), with both being more severe in the class with less profound methylation changes. The lack of correspondence between molecular signatures, including methylation and gene expression, and clinical manifestations caution against making phenotypic inferences based on the blood-based methylation profile. This is important to consider when evaluating the use of these episignatures as biomarkers for patient stratification and/or response to pharmacological agents in clinical trials. In the specific case of ADNP syndrome, these observations are timely as potential experimental therapeutics emerge. In particular, pre-clinical data suggest that an eight amino acid ADNP peptide called NAP (NAPVSIPQ) might hold promises for treatment. NAP has been shown to have broad neuroprotective effects *in vitro* and *in vivo* ^35^, and *in vivo* administration of NAP or its derivatives in a mouse model of ADNP syndrome has been shown to ameliorate synaptic and behavioral defects ^10–13^. In conclusion, while the episignatures in ADNP syndrome have proven very valuable for the purpose of complementing clinical genetics for the purpose of enhancing accuracy of diagnosis, their value as biomarkers in clinical trials should be carefully evaluated.

## Acknowledgements

This work was supported by grants from the Beatrice and Samuel A. Seaver Foundation and the ADNP Kids Research Foundation. The work was also supported by the Spanish Ministry of Economy, Industry and Competitiveness, and Instituto de Salud Carlos III. Research reported in this paper was supported by the Office of Research Infrastructure of the National Institutes of Health under award number S10OD018522. This work was supported in part through the computational resources and staff expertise provide by Scientific Computing at the Icahn School of Medicine at Mount Sinai.

## Figure Legends

**Table S1.** Genetic and demographic information about the 43 individuals included in the study. For each patient, the position of the genetic mutation on cDNA, protein and genomic DNA, the exon location, the effect, the inheritance, the classification of pathogenicity, and the mutation class are shown. Demographic information includes sex and country of origin.

**Table S2.** Differentially methylated CpGs and differentially expressed genes in individuals with class I and class II *ADNP* mutations.

**Table S3.** Extended version of Table 1, with separate and combined analyses of cohorts S and W across all clinical measures.

**Figure S1.**
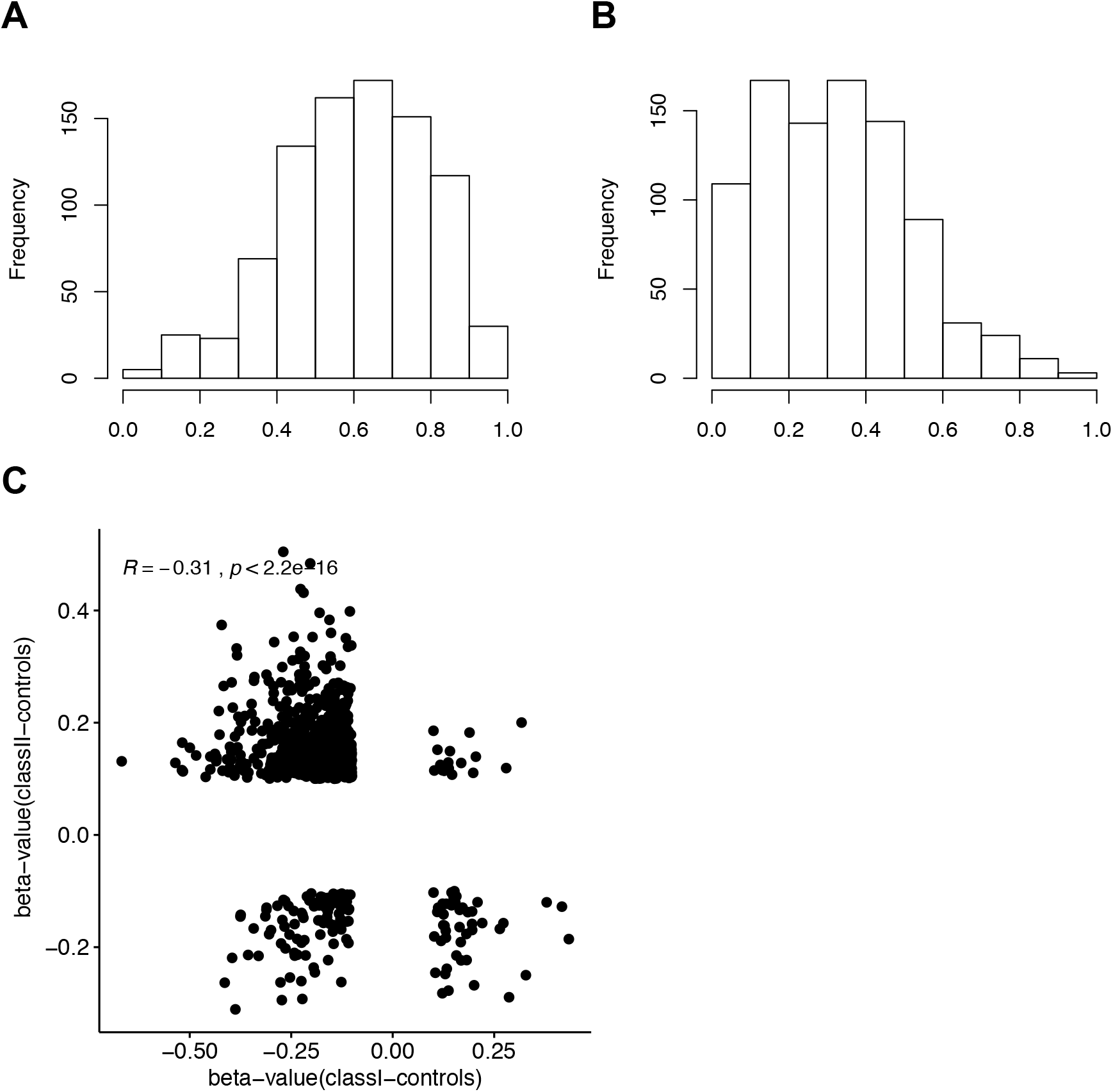
**(A)** Distribution of the mean β-values for the 888 differentially methylated CpGs shared between the two classes for individuals carrying class I mutations. **(B)** Distribution of the mean β-values for the 888 differentially methylated CpGs shared between the two classes for individuals carrying class II mutations. **(C)** Correlation between (β -values_cases_-β -values_controls_) in class I (X-axis) and (β -values_cases_-β -values_controls_) in class II (Y-axis).

**Figure S2.**
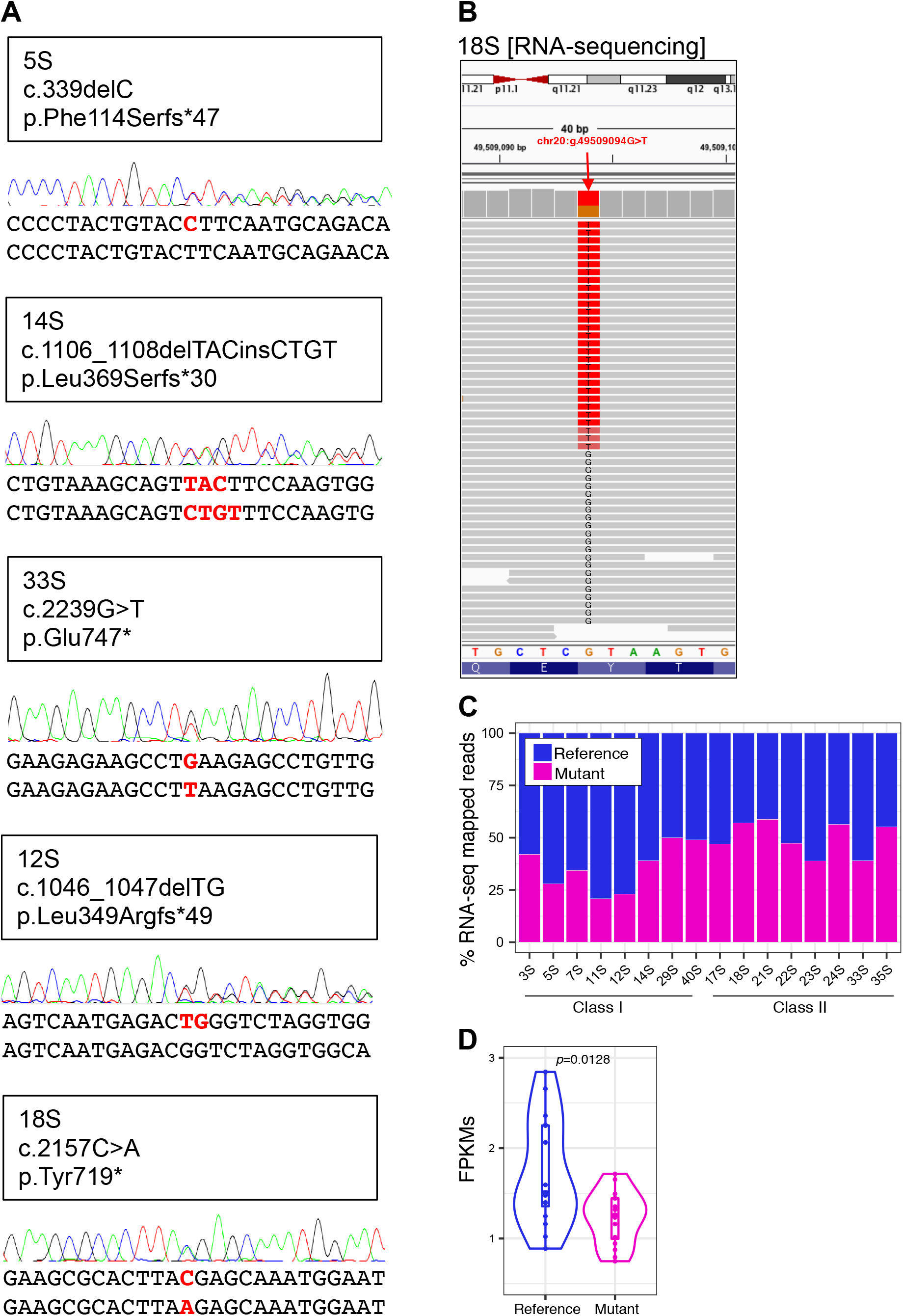
Biallelic expression of *ADNP* mutant mRNAs for (**A**) three class I mutations (p.Phe114Serfs*47, p.Leu349Argfs*49, and pLeu369Serfs*30) and two class II mutations (p.Tyr719 and p.Glu747*). The Sanger sequencing traces, alongside the cDNA sequence for reference (top) and mutant (bottom) alleles are shown. (**B**) RNA-sequencing pile-up of class II mutation p.Tyr719 (sample 18S) displaying coverage and number of mutant reads (red) and reference (*i.e.* healthy) reads (grey). (**C**) RNA-sequencing quantified the total fraction of *ADNP* mutant alleles and reference alleles present for each sample. Note, sample 1S was not included in this analysis (Table S1). (**D**) The distribution of expression (FPKM) for all mutated reads aggregated together relative to reference reads of the *ADNP* gene across all samples. We observed lower expression of the mutant allele (Mann-Whitney U Test, *p*=0.01).

**Figure S3.**
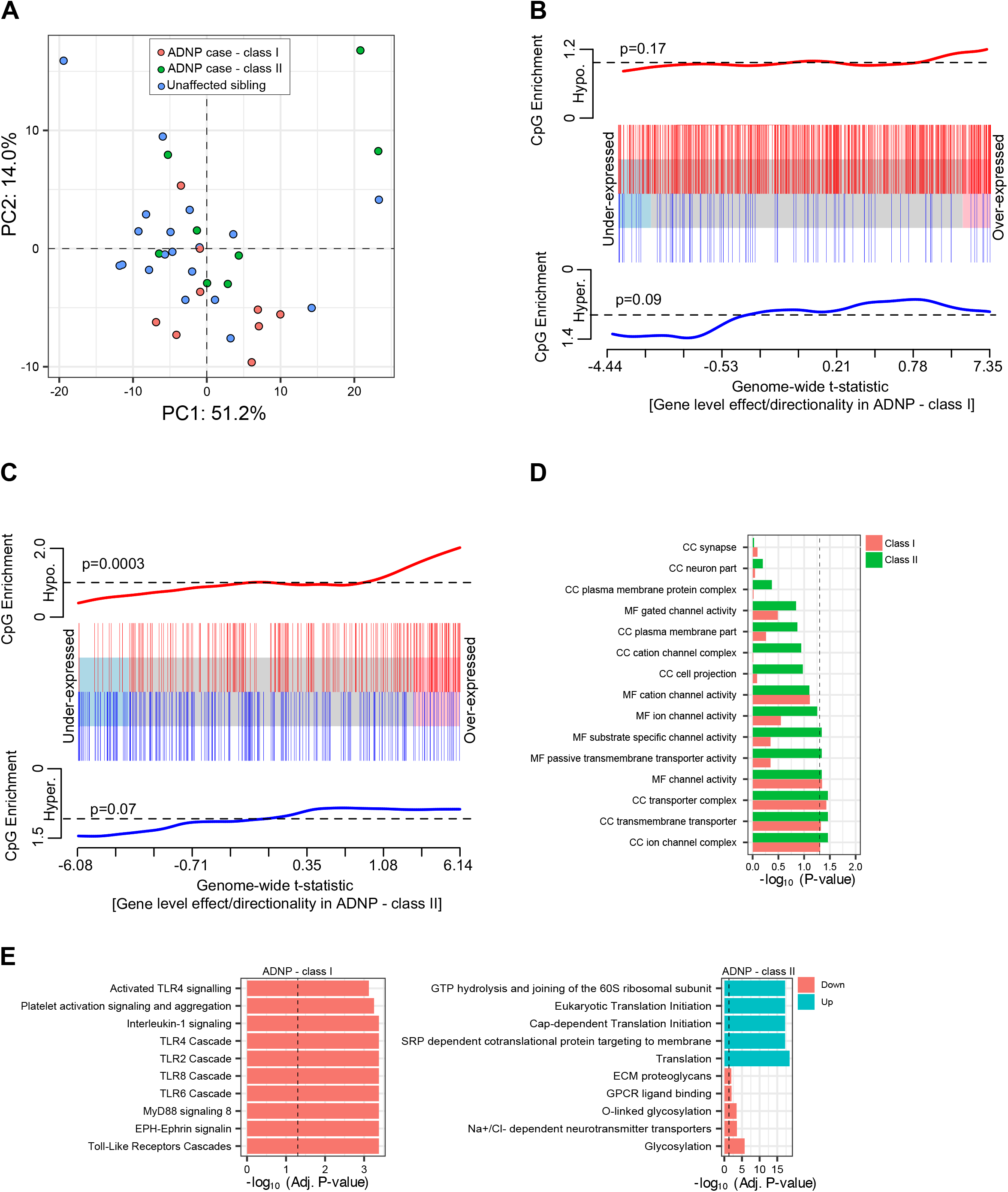
(**A**) Principal component analysis (PCA) on the RNAseq data from 17 ADNP cases (class I, red; class II green) and 19 unaffected sibling controls (blue), showing no distinct separation between mutational classes or controls. PCA was constructed using only genes harboring significant differentially methylated CpGs within their respective gene body reported in Fig 1A-B. Competitive gene-set enrichment analysis examined over- and under-expression of genes harboring significantly hypomethylated and hypermethylated CpGs, respectively for (**B**) ADNP cases with class I mutations and (**C**) and ADNP cases with class II mutations. Hypomethyated CpGs mapped to over-expressed genes in ADNP cases with class II mutations (p=0.0003). No other significant associations were detected. (**D**) RNA-seq supervised GO enrichment of pathways found to be significantly enriched for differentially methylated CpGs. (**E**) Exploratory GO enrichment analysis and the top ten under-expressed pathways in ADNP cases with class I mutations (left) and top 5 under-expressed and top 5 over-expressed pathways ADNP cases with class II mutations (right). No pathways were over-expressed in ADNP cases with class I mutations.

