## Supplemental Note (extended materials and methods) for "Episignatures stratifying ADNP syndrome show modest correlation with phenotype"

**Supplemental Subjects and Methods**

**Cohorts, patients and genetic information**

The study comprises four cohorts. Cohort A was collected as part of the Autism Sequencing Consortium ^1; 2^. Cohort R was collected at the Medical Genetics Laboratory of Ospedale San Camillo-Forlanini ^3^ or Ospedale Pediatrico Bambino Gesù, Rome, Italy. Cohort S was collected at the Seaver Autism Center for Research and Treatment, Icahn School of Medicine. Cohort W was collected at the University of Washington. Mutations in Cohort A were identified through research-based whole-exome sequencing ^1; 2^ and validated by Sanger sequencing. Mutations in Cohort R, S, and W were identified and validated in Clinical Laboratory Improvement Amendments (CLIA)-certified laboratories. Participation was approved by the Institutional Review Boards of participating sites. All caregivers provided informed written consent and assent was obtained when appropriate. We used 24 cases (cohorts A, R, and part of S) for methylation analyses; 30 cases (S and W) for phenotypic analyses; and, 17 cases (part of cohort S) for RNA-sequencing analysis. The 19 unaffected age-matched and 14 unaffected sibling controls were from cohort S. All mutations are described according to the Human Genome Variation Society (HGVS) guidelines for mutation nomenclature. The cDNA and amino acid positions are annotated according to the most updated *ADNP* RefSeq mRNA and protein sequence (NM_015339.4 and NP_056154.1). Nucleotide numbering referring to cDNA uses +1 as the A of the ATG translation initiation codon in the reference sequence, with the initiation codon as codon 1. Further, to generate the lollipop plot in **Figure 1A**, we searched the published and re-annotated literature for mutations in *ADNP* using PubMed, HGMD Professional (Biobase), and denovo-db. We also included pathogenic and likely pathogenic mutations deposited in ClinVar (NCBI, <http://www.ncbi.nlm.nih.gov/clinvar/>).

**Clinical evaluation**

Prospective clinical and psychological characterization was completed for 22 individuals seen at the Seaver Autism Center and 10 individuals seen at the University of Washington. A battery of standardized assessments was used to examine ASD, intellectual functioning, adaptive behavior, language, motor skills, and sensory processing (see below). The medical evaluation included psychiatric, neurological, and clinical genetics examinations, and medical record review.

*ASD phenotype.* Gold-standard ASD diagnostic testing included the Autism Diagnostic Observation Schedule, Second Edition (ADOS-2) ^4^, the Autism Diagnostic Interview-Revised (ADI-R) ^5^, and a clinical evaluation to assess Diagnostic and Statistical Manual for Mental Disorders, Fifth Edition (DSM-5) criteria for ASD ^6^. A consensus diagnosis was determined based on results from the ADOS-2, ADI-R and the clinical evaluation. The ADOS-2 and ADI-R were administered and scored by research reliable raters and the psychiatric evaluation was completed by a board-certified child and adolescent psychiatrist or licensed psychologist. The ADOS-2 is a semi-structured observational assessment that provides scores in the domains of Social Affect, Restricted and Repetitive Behavior, and a total score. A comparison score ranging from 1-10, with higher scores reflecting a greater number of symptoms, was calculated to examine symptom severity within each ADOS-2 domain and in total ^7^. 21 individuals received Module 1 of the ADOS, for children who are nonverbal or communicate using single words. Eight individuals received Module 2, for individuals who communicate using phrase speech. One individual received Module 3, for children who are verbally fluent. The ADI-R is a structured caregiver interview that assesses ASD symptomatology within the domains of socialization, communication, and repetitive and restricted interests and behavior. A consensus diagnosis was determined for each participant based on results from the ADOS-2, ADI-R and clinical evaluation using DSM-5.

*Intellectual functioning.* Global cognitive ability was measured using the Mullen Scales of Early Learning ^8^, the Stanford Binet Intelligence Scales, Fifth Edition ^9^, or the Differential Ability Scales, Second Edition (DAS-II) ^10^, depending on age and verbal ability. The Mullen is validated for children from birth to 68 months, but is commonly used for older individuals with ID ^11^. Developmental quotients were calculated using age equivalents divided by chronological age as has been done in previous studies ^12^. For example, a nonverbal developmental quotient was computed by dividing the mean age equivalents on the visual reception and fine motor scales by the child’s chronological age and then multiplying by 100. The DAS-II is a measure of cognitive functioning that assesses a child’s verbal reasoning, nonverbal reasoning, and spatial abilities. A general conceptual ability index can be calculated to assess overall intellectual functioning. The Stanford-Binet Intelligence Scales, Fifth Edition is an intelligence test that produces a nonverbal intellectual quotient (IQ), verbal IQ, and full scale IQ based on performance across five scales: fluid reasoning, knowledge, quantitative reasoning, visual-spatial, and working memory.

*Adaptive behavior.* The Vineland Adaptive Behavior Scales, Second Edition, Survey Interview Form (Vineland-II) ^13^ and the Vineland Adaptive Behavior Scales, Third Edition, Comprehensive Interview Form (Vineland-3) are clinician-administered interviews that assesses adaptive behavior in the domains of communication, daily living skills, socialization, and motor skills. The Vineland-II was completed for 13 individuals. The Vineland-3 was completed for 19 individuals. The motor domain is intended for children ages six years and under, but was assessed in 19 individuals given significant motor delays in this population. The Vineland-II and Vineland-3 were also used in conjunction with cognitive testing to identify the presence and severity of ID.

*Language skills*. Language milestones were assessed during the ADI-R and the psychiatric evaluation. Current expressive and receptive language abilities were assessed using the Mullen, Vineland-II, MacArthur-Bates Communicative Development Inventories ^14^, Peabody Picture Vocabulary Test, Fourth Edition ^15^, and Expressive Vocabulary Test ^16^.

*Motor skills.* Motor milestones were assessed during the ADI-R and the psychiatric evaluation. Current motor skills were assessed using the Vineland-II and Mullen fine and gross motor skills domains. The Beery Visual-Motor Integration Test, 6^th^ Edition ^17^ was completed when appropriate.

**Genome-wide methylation arrays and data quality control**

For the methylation analyses, genomic DNA was isolated from the peripheral blood of 24 ADNP cases, 19 unaffected age-matched, and 14 unaffected siblings from cohorts S, A and R. DNA methylation analysis was performed using the Ilumina EPIC 850K methylation arrays, according to the manufacturer’s protocol.

The sex of the individuals included in the study was inferred both by the number of probes on chromosome Y (chrY) with detection p>0.01, and the mean β value of probes on chromosome X (chrX) per sample. Samples with high failure rate for chrY probes and high mean β value for chrX probes were inferred as females, while samples with low failure rate for chrY probes and low mean β value for chrX probes were inferred as males. Epigenetic age was predicted using the online tool published by Horvath, 2013 ^18^. Where chronological age at sampling was recorded, the mean discordance between predicted age and observed age for 24 samples was +/- 1.9 years, which is similar to previous reports ^18^. The fraction of circulating peripheral blood cell types were predicted for each sample using the method of Houseman et al. 2012 ^19^. Predications for the following cell types were obtained: CD4+ T cells, CD8+ T cells, natural killer cells, B lymphocytes, monocytes and granulocytes.

Data from *ADNP* cases and controls were normalized as described in previous studies ^20; 21^. In short, 862,927 probe sequences (50-mer oligonucleotides) were remapped to the reference human genome hg19 (NCBI37) using BSMAP, allowing up to 2 mismatches and 3 gaps, to retain uniquely mapped autosomal probes. We removed any probe that overlapped SNPs with MAF ≥5% identified by the 1000 Genomes Project within 5 bp upstream of the targeted CpG. Further, we removed probes with a detection p>0.01 in each individual. After filtering, we retained 820,167 autosomal probes, which were subjected to background correction, two color channel normalization and quantile normalization using the *lumi* package in R ^22^ The distributions of Infinium I and Infinium II probes were adjusted using *BMIQ* ^23^*.* Probes were then annotated based on their position relative to RefSeq genes using BEDTools v2.17 ^24^. We defined promoter regions as ±2 kb from transcriptional start sites, gene body regions as transcription start to transcription end, and intergenic regions not annotated by the preceding categories. We also annotated individual CpG based on their overlap with CpG islands based on annotations in the UCSC genome browser, CpG shore (±2 kb of island), CpG shelf (±2 kb of shore), and CpG sea (regions outside the previous three categories).

**Identification of an episignature in *ADNP* cases**

We performed linear regression using age, sex, predicted blood cell composition and *ADNP* mutation status as independent variables (Test Model: Methylation ~ Disease status + Age + Gender + CD4T + natural killer cells + B cells + Monocytes + Granulocytes + Batch). Regression analysis was completed separately for class I and class II *ADNP* mutations. We did not include CD8T cell composition in the model due to very low abundance across all samples. We selected probes associated with disease status at 1% FDR and with minimum β value difference between ADNP cases and controls ≥0.1. Principal component analysis and unsupervised clustering of episignatures was performed on methylation data following Combat batch adjustment ^25^ to remove systematic sources of variability related to batches without introducing false signal.

**Enrichment analyses**

We assessed enrichment of this set of genes with differentially methylated CpG using ing gene lists: *a)* 914 genes implicated in developmental delay/intellectual disability (DD/ID), based on the Developmental Disorders Genotype-Phenotype Database (DDG2P) (<https://decipher.sanger.ac.uk/info/ddg2p>), *b)* 102 ASD risk genes defined by the Autism Sequencing Consortium ^2^, and, *c)* 145 risk genes for CHD build in house. In each test, we utilized a background gene set comprising 25,141 genes that overlapped 606,904 probes covered by the Illumina 850K EPIC methylation arrays. For each comparison between the differentially methylated genes in the input set, we first constructed the empirical distribution by randomly sampling the same number of genes as in the input set from the background gene set 10,000 times, using a custom script in R. Enrichment p-values were computed by calculating the number of sampled gene lists that had at least as many overlapping genes with the target sets as the input set, divided by 10,000 permutations.

**RNA isolation, library preparation, and quantification of gene expression**

Blood was collected for 17 ADNP cases and 19 unaffected siblings using PAXgene RNA tubes (Qiagen, Valencia, CA, USA) and total RNA was extracted and purified in accordance with the PAX gene RNA kit per manufacturer’s instructions. Globin mRNA was depleted from samples using the GLOBINclear Human Kit (Life Technologies, Carlsbad, CA, USA). The quantity of purified RNA was measured on a Nanodrop 2000 Spectrophotomerter (Thermo Scientific; 61.4 ± 24.1 ng μl^−1^) and RNA integrity numbers (RIN) measured with the Agilent 2100 Bioanalyzer (Agilent, Santa Clara, CA, USA; 8.31 ± 0.68). The Illumina TruSeq Total RNA kit (Illumina, San Diego, CA, USA) was used for library preparation accordingly to manufacturer instructions without any modifications. The 36 indexed RNA libraries were pooled and sequenced using long paired-end chemistry (2x150 bp) at an average read depth of 10M reads per sample using the Illumina HiSeq2500. Illumina adapter sequences were trimmed from all fragmented reads using TrimGalore (options –paired –illumina) (https://www.bioinformatics.babraham.ac.uk/projects/trim_galore/). All high-quality trimmed reads were mapped to UCSC *Homo sapiens* reference genome (build hg37) using default STAR v2.4.0 parameters ^26^. Samtools was used to convert bamfiles to samfiles and featureCounts ^27^ was used to quantify gene expression levels for each individual sample using default paired-end parameters. RNA-sequencing FASTQ files are available at the Gene Expression Omnibus under accession number GSEXXXXX (accession number will be updated post publication).

**RNA-seq data quality control**

Raw count data measured 56,632 genes across 36 subjects. Unspecific filtering removed lowly expressed genes that did not meet the requirement of a minimum of 1 count per million (cpm) in at least 8 subjects (40% of subjects). A total of 20,491 genes were retained, then subjected to edgeR VOOM normalization ^28^, a variance-stabilization transformation method. Normalized data were inspected for outlying samples using unsupervised hierarchical clustering of subjects (based on Pearson coefficient and average distance metric) and principal component analysis to identify potential outliers outside two standard deviations from these averages. No outliers were present in these data.

**Mutant allele abundance**

We took measures to reduce the likelihood of false positives and biases in quantification of relative allele abundances at each mutation location. RNA-seq reads were filtered and mapped to retain only uniquely mapped reads, using STAR ^24^ and SAMtools, and then further removed potential PCR duplicates using the MarkDuplicates function in Picard tools with default parameters (http://broadinstitute.github.io/picard/faq.html). Allelic counts were computed at each single-nucleotide variant position in each sample using the SAMtools mpileup function. Next, we recorded the number of overlapping sequences containing the mutant and reference allele. The mutant allele frequency was defined as the number of covering RNA-seq reads containing the mutant allele at that position, divided by the total number of RNA-seq reads overlapping that position.

**Differential gene expression and functional annotation analyses**

A moderated *t*-test, implemented through the *limma* package ^28^, assessed differential gene expression between unaffected siblings and *ADNP* cases with class I and class II mutations, respectively. The analysis covaried for the possible influence of sex, gender, RIN and sequencing batch on gene expression differences. Significance threshold was set to a Benjamini-Hochberg multiple test corrected *P*-value <0.05. Correlation adjusted mean rank (CAMERA) gene set enrichment ^29^ was performed using the two sets of resulting summary statistics for ADNP cases with class I and class II mutations. CAMERA performs a competitive gene set rank test to assess whether the genes in a given set are highly ranked in terms of differential expression relative to genes that are not in the set. The test ranks gene expression differences in ADNP cases with class I and class II mutations relative to unaffected siblings, respectively, to test whether gene sets are over-represented towards the extreme ends of these ranked lists. It uses ***limma***’s linear model framework and accommodates the observational-level weights from voom in the testing procedure. After adjusting the variance of the resulting gene set test statistic by a variance inflation factor that depends on the gene-wise correlation (which we set to default parameters, 0.01) and the size of the set, a p-value is returned and adjusted for multiple testing. This approach was used to examine whether changes in gene expression were observed for genes harboring significant differentially methylated CpGs in ADNP cases and to explore changes in REACTOME pathways.

**RT-PCR and Sanger sequencing**

Reverse transcription was performed on 50ng of RNA using High Capacity cDNA Reverse Transcription Kit (Applied Biosystems, California, USA) according to the manufacturer’s protocol. PCR was performed using FastStart PCR Master (Roche, Mannheim, Germany) or Phusion High-Fidelity PCR Kit (New England BioLabs, Ipswich, Massachusetts). A specific primer spanning the exon 2-exon 3 junction of ADNP with sequence 5’-ACGAAAAACCAGGACTATCGGA-3’ was used as the forward primer in all PCR reactions. Reverse primers were mutation-specific and as follows: 5S, 5’-AAACAGCTTGCTCTACACTGTCA-3’; 12S 5’- TGGCCCGATGAGAGAGAAGA-3’; 14S, 5’-CTGCAGCAGGTTTGGAACTG-3’; 18S, 5’-TGGTGGGATAGGGCTGTTTG-3’; and, 33S, 5’- TAAACTGGCTGCTAGCTTCTCAA-3’. PCR products were purified using a MultiScreen PCR filter plate (Millipore Sigma, Burlington, Massachusetts) and submitted for Sanger Sequencing at Genewiz with the same reverse primers used in the PCR reaction and forward primers as follows: 5S 5’- ACGAAAAACCAGGACTATCGGA-3’; 12S 5’-ATCGGTTCCCTTGCTTCTGG -3’; 14S 5’- TGCAGCAGAACAACTATGGAGT-3’; 18S 5’-ATACCAGCAACATGACCGCC-3’; 33S 5’- CACCCTCTCGGCTTAATCAGT-3’.
